# HIV-1 GAG SPECIFICITY FOR PIP_2_-CONTAINING MEMBRANES MIGHT BE DRIVEN BY MACROMOLECULAR ELECTRIC PROPERTIES RATHER THAN MOLECULAR AFFINITIES

**DOI:** 10.1101/2022.08.25.505363

**Authors:** L. B. P. Socas, E. E. Ambroggio

## Abstract

The HIV-1 assembly occurs at the plasma membrane, where the GAG polyprotein plays a crucial role. The GAG-membrane association is directed by the matrix domain (MA), which is myristoylated and has a highly basic region that interacts with the anionic lipids. Several evidence suggests that the presence of phosphatidylinositol-(4,5)-bisphosphate (PIP_2_) highly influence this binding. In addition, MA also interacts with nucleic acids, which is proposed to be important for the specificity of GAG for PIP_2_-containing membranes. It is proposed that RNA could have a chaperone function when interacting with the MA domain, preventing GAG from associating with unspecific lipid interfaces. Here, we study the interaction of MA with monolayer and bilayer membrane systems, focused on the specificity for PIP_2_ and on the possible effects of a GAG N-terminal peptide to impair the binding for either RNA or membrane. We found that RNA decreases the kinetics of the protein association with lipid monolayers but without any effect on the selectivity for PIP_2_. Interestingly, for bilayer systems, this selectivity increases in presence of both the peptide and RNA, even for highly negative charged compositions, where MA by itself doesn’t discriminate between membranes with or without PIP_2_. Therefore, we propose that the specificity of MA for PIP_2_-membranes might be related to the electrostatic properties of both membrane and protein local environments, rather than a simple difference in molecular affinities. This scenario gives a new understanding of the regulation mechanism with a macromolecular view instead of considering molecular interactions within a ligand-receptor model.

**Importance:** HIV-1 virions are formed at the PM of infected cells through a direct interaction of the viral GAG protein with lipids. This is a finely regulated process governed by the GAG N-terminal matrix domain, MA. Here, we obtained compelling evidence on how this process depends on the local dielectric environments of both, the membrane and MA. Using bio-membrane mimicking systems, we found how MA myristoylation is involved in the interfacial absorption and anchoring of the protein, where the interaction with RNA negatively regulates this process but in a lesser extent when traces of the PIP_2_ lipid are present. Additionally, an N-terminal GAG-derived peptide competes with MA for the nucleic acid binding and impair the protein-membrane interaction when PIP_2_ is absent. All these data allowed us to propose a model for MA association with lipid interfaces and how it depends on oligonucleotide binding, lipid composition and competing peptide presence.

## 1. Introduction

The second stage of the HIV-1 lifecycle is characterized by the production of structural proteins, the assembly of viral particles and their subsequent exit from the host plasma membrane (PM), generating new viruses ready to infect other cells (1-3). The immature virus formation is proposed to directly occur at the PM, where the GAG polyprotein plays a crucial role in coordinating such a complex process (4-11). The association of GAG with the membrane is directed by its matrix domain (MA), located in the amino terminal region (10-16). This domain is myristoylated and has a highly basic region (HBR) responsible for the interaction with the negatively charged lipids of the membrane (8, 11, 16-18). There is compelling experimental evidence suggesting that the binding of GAG or MA to membranes is an electrostatic driven process highly influenced by the presence of phosphatidylinositol-(4,5)-bisphosphate (PIP_2_) (6, 15, 17-19). This lipid is proposed as some sort of receptor that, together with the exposure of the protein’s myristic group, helps with the anchoring of GAG to the PM (19-21).

GAG polyprotein is also known to bind nucleic acid molecules, including not only the viral genome, but also certain oligonucleotides and microRNAs (22-28). These bindings could involve both the nucleocapsid and matrix domain of GAG. Purohit et al. in 2001, by using the SELEX technique, identified a consensus RNA sequence that specifically binds to the MA domain with high affinity. They also showed that this protein-RNA interaction is mediated by a group of basic residues in MA (23). This 13-bases consensus sequence, flanked on both sides by nucleotides forming a duplex, is known as Sel25 and is widely used as a model for studies of MA binding to nucleic acids (23, 28). More recent findings have shown that, in the cellular context, the MA domain binds almost exclusively, and very selectively, to certain tRNAs (25, 26). In fact, Kutluay and colleagues in 2014 noted that the most frequent binding event between GAG and RNA in cells was the binding of MA to tRNA (25). In this same line, Gaines et al. in 2018 studied the interaction of MA with tRNA^LYS3^, an isoform of tRNA of particular interest due to its role as an initiator in the reverse transcription process. Using NMR experiments, they confirm that tRNA^LYS3^ interacts with the same group of basic residues important for the PIP_2_ binding (26).

This MA-RNA interaction is proposed to be important for the specificity of GAG for PIP_2_-containing membranes (17, 27-30). Several studies have shown that the binding of MA to nucleic acids competes for the protein interaction with negatively charged membranes, which can be overcome by the presence of PIP_2_ (27, 28). In addition, RNase treatment of GAG extracts obtained from mouse reticulocytes reduces the selectivity of the protein for PIP_2_-containing liposomes (30). All this evidence suggests that the RNA could have a chaperone function when interacting with the MA domain, preventing GAG from associating with lipid interfaces until it reaches the PIP_2_-rich PM (9, 17, 28, 31). Several authors have proposed that this would be plausible if the affinity of MA for nucleic acids were lower than its affinity for PIP_2_ and higher than for other phospholipids, implicating that this interaction would protect the protein from binding to inappropriate membranes (27, 28). Therefore, finding ways to impair with this regulatory mechanism could be useful to properly understand viral assembly, in order to be able to develop potential therapies that target this stage of the virus lifecycle.

In a previous works, we studied the properties of a peptide derived from the N-terminal region of GAG, comprising the first 21 amino acids of the protein (myrMA_21_), which contains part of the HBR region and the myristic group (32, 33). We found that this peptide is able to interact whit nucleic acids with high stoichiometry due to its formation of aggregates in solution (32). Interestingly, this peptide is able to partition into negatively charged membranes but doesn’t show any selectivity toward PIP_2_, probably due to the lack of the complete MA domain sequence (33). Here, we study the interaction of MA with membranes, using both monolayer and bilayer systems, focused on the specificity for PIP_2_ and on the possible competition of myrMA_21_ for the binding of either RNA or membrane. In this sense, we found that RNA decreases the kinetics of the protein adsorption toward the lipid interface in a monolayer system but without any effect on the selectivity for PIP_2_. On the other hand, the peptide competes very strongly for the binding of RNA, however, this competition is not enough to overcome the oligonucleotide influence on the MA specificity for PIP_2_-containing membranes in a bilayer system. Interestingly, we show that the presence of the peptide and/or RNA increases the selectivity of MA for PIP_2_, even for highly negative charged compositions, where MA by itself doesn’t discriminate between membranes lacking or containing PIP_2_. Considering these and previous results, we believe that the regulation of the specificity of MA for PIP_2_-containing membranes might be related to a finely tuned dielectric properties of both membrane and protein local environment and not the result of different molecular affinities between protein-nucleic acids and protein-membranes.

## 2. Materials and methods

### 2.1. Materials and reagents

The peptide myrMA_21_ [myr-GARASVLSGGELDKWEKIRLR-NH2] was purchased from Biomatik USA, LLC (USA, DE). The samples used for the experiments were prepared diluting the peptides in acetonitrile 30% (as recommended by the provider) and the final concentration was determined by UV absorption spectroscopy (ε_280nm_ = 5690 M^-1^cm^-1^). The purity of the peptide, judged by HPLC, was greater than 95%. The RNA Sel25 oligonucleotide (5’GGACAGGAAUUAAUAGUAGCUGUCC3’) was obtained from Invitrogen (USA) along with a FITC tagged version on the 5’ extreme. For the experiments presented in this work, the oligonucleotide was diluted in DNase-free and RNase-free water.

Lipids 1-palmitoyl-2-oleoyl-sn-glycero-3-phosphocholine (POPC); 1,2-dioleoyl-sn-glycero-3-phospho-L-serine (sodium salt) (DOPS); L-α-phosphatidylinositol-4,5-bisphosphate (Brain, Porcine) (ammonium salt) (PIP_2_) were purchased from Avanti Polar Lipids, Inc. (Alabama, U.S.A.); while 1,1′-dioctadecyl-3,3,3′,3′-tetramethylindodicarbocyanine-5,5′-disulfonic acid (DilC_18_) was purchased from Invitrogen. All stock solutions were prepared in chloroform/methanol (2:1, v/v) except for the PIP_2_ case, where a 4% of water was included.

### 2.2. Protein purification

Non-myristoylated HIV-1 MA as well as the myristoylated version (myrMA), were expressed in *Escherichia coli* strain BL21(DE3) transformed with pET-11a-based vectors kindly provided by Dr. Michael F. Summers (University of Maryland Baltimore County) as described in previous works (32, 34). The myristoylated variant was co-expressed along with the *Saccharomyces cerevisiae* N-myristoyltransferase and, in this case, myristic acid was included in the bacterial growth media during the induction phase. Both proteins contain a 6His tag in order to use a Ni-NTA affinity chromatography as a capture step, which was followed with an intermediate purification by either an SP-HiTrap cation exchange chromatography for MA or a butyl-HiTrap hydrophobic interaction chromatography for myrMA. Before each experiment, a final purification step by molecular exclusion chromatography was performed using a Superdex 75 HR column equilibrated with 20 mM Tris-HCl (pH 7.4) 150 mM NaCl.

### 2.3. Liposome preparation

For liposome preparation, the appropriate moles of lipids were mixed in conic glass tubes, slowly dried with a steam of nitrogen gas and left in vacuum for 2 – 3 hours. These dried lipid films were resuspended in 20 mM Tris-HCl (pH 7.4) 150 mM NaCl. Large unilamellar vesicles (LUVs) were prepared by freeze–thaw and extrusion (35) in a device from Avestin (Ottawa, Canada). Three cycles of freeze and thaw were done using N_2_ (l) and the samples were extruded 20 times through a 100 nm pore size polycarbonate membrane from Whatman (Schleicher & Schuel) at room temperature. Each sample were prepared to a final lipid concentration of 4 mM and used for experiments within 3 days. After each preparation, liposome samples were checked by dynamic light scattering with a Submicron Particle Sizer equipment (Nicomp 380, Santa Barbara, CA) and in all cases a monodisperse population was observed, with a mean diameter size of around 110 nm (variation: ± 5%).

### 2.4. Monolayer studies

The monolayer studies were done on a control unit Monofilmmeter (with Film Lift, Mayer Feintechnique, Germany). The surface pressure (Π) was measured using the Wilhelmy method with a Platinized-Pt plate and the data were continuously registered with a double channel X-YY recorder. The monolayers were formed by direct spreading the lipids from chloroform/methanol (2:1, v/v) solution using a micro syringe and the solvent was allowed to evaporate for at least 5 min before the measurements. The adsorption and penetration assays were done by injecting the proteins from their aqueous solution into 15 mL of subphase contained in a 15 cm^2^ trough. For the mixing studies, monolayers were formed by spreading the protein directly onto the previously constructed lipid interface at different protein-lipid proportions. In this case, the total surface area range of the Langmuir trough was 12 - 80 cm^2^ and the isothermal compression were carried out at a fixed rate of ∼0.0475 nm^2^·molecule^-1^·s^-1^. For the effect of the RNA, the protein samples were pre-incubated with a 5-fold amount of the Sel25 oligonucleotide. All the experiments were performed at room temperature, with a subphase containing Tris 20 mM (pH 7.4) and NaCl 150 mM and independently repeated at least 3 times.

For the analysis of the adsorption and penetration experiments, information was empirically obtained by fitting an exponential equation to the Π-time data (36):

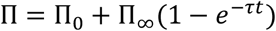

Where Π_0_ and Π_∞_ indicate the initial and final surface pressure (value at *t* → ∞) respectively, and *τ* denotes the time constant of adsorption of the protein toward the interface. From *τ* is possible to obtain the interfacial rate of adsorption (*κ*) as the first derivative evaluated at *t* = 0, i.e., κ = Π_∞_*τ*.

In addition, the thermodynamic tendency of the protein to partition into a lipid-free interface can be estimated by calculating the free energy of adsorption, evaluated by (37):

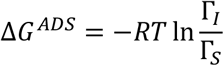

where Γ_*I*_ and Γ_*S*_ are the molar concentrations of the protein at the interface and subphase, respectively, T is the temperature (295 K), and R is the gas constant (1,987 cal·mol^-1^·K^-1^). Γ_*I*_ is calculated from the molecular area obtained via the compression isotherm of pure protein monolayer, considering a total surface area of 15 cm^2^.

For the miscibility analysis we used the Λ-factor, a previously proposed methodology for comparing compression isotherms (32). Briefly, the Λ-factor is defined as the pair (λ, r^2^), where λ quantifies the area shift of one isotherm with respect to the other and r^2^ denotes the similarity between the curves. In this sense, we used this method to compare the experimental isotherm obtained from lateral compression of the protein-lipid mixed film with respect to the theoretical ideal one, constructed by using the following expression:

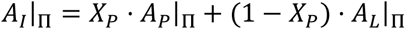

Were *A*_*I*_ is the mean molecular area for the ideal case at pressure Π, *A*_*P*_ and *A*_*L*_ are respectively the areas of the pure protein and lipid films at pressure Π and *X*_*P*_ is the protein molecular fraction.

The calculated ideal isotherm represents a protein-lipid composed film with no net lateral interactions, therefore, deviations from this behavior toward higher or lower areas will represent the existence of repulsive or attractive interactions, respectively (37-41). The Supporting Information section S1 contains an explanation of the Λ-factor.

### 2.5. Beads binding assay

In order to quantify the binding of MA to liposomes and/or oligonucleotides we used a previously developed binding assay based on affinity chromatography beads (27). Briefly, agarose beads modified with Ni-NTA (Qiagen), were equilibrated with a washing solution (20 mM Tris-HCl, pH 7.4, 150 mM NaCl) and incubated with 6His tagged MA protein for 2 hours at 4°C. Afterward, the packed beads were pelleted, washed twice with 300 μL of washing solution, and resuspended with a total volume of 500 μL of the same solution. These beads with immobilized protein were kept at 4ºC for no more than three days prior to use.

For the binding assays, 20 μL of the protein-loaded beads suspension were incubated with 50 μL of the mix of interest that, depending on the experiment, contained liposomes labeled with 0.5% of the fluorescent DilC_18_ lipid (1 mM of total lipids) with or without myrMA_21_ and/or 10 μM of Sel25. In the case of the experiments where binding to oligonucleotides was observed, the same amount of a FITC labelled variant of Sel25 was used. The mixture was incubated for at least one hour at room temperature, after which the beads were sedimented by centrifugation (1 min at 5000 g), rinsed twice with 300 μL of washing solution and resuspended in 50 μL of the same solution. After this procedure, the samples were observed under an epifluorescence microscope.

### 2.6. Microscopy measure and quantification

For the beads visualization and binding quantification, a Zeiss AxioPlan fluorescence microscope was used with a 20X LD A-Plan objective and equipped with an XM10 monochrome camera with the CellSens Entry program (Olympus) for image acquisition. The samples to be visualized were placed in microcells constructed between a slide and a coverslip separated by 1 mm with Parafilm tape and with a volume capacity of approximately 20 μL. Images were taken using a Zeiss Set 10 filter for green (excitation: 450 – 490 nm; dichroic: FT 510; emission: 515 – 565 BP) and a Zeiss Set 15 filter for red (excitation: 546/12 BP; dichroic: FT 580; emission: 590 LP). For each individual experiment, exposure and gain values were adjusted identically. The images were analyzed using ImageJ software, for which the fluorescence intensity of the region of interest was quantified. For the calculation of the relative brightness (intensity ratio between a sample and a reference), all the conditions used as reference were measured in parallel to the samples on the same day. All measurements were performed at least three times independently. For each case, control measurements were made using beads without immobilized protein, where no significant non-specific binding was observed (data not shown).

## 3. Results

### 3.1. MA interaction with lipid monolayers

The interaction of GAG with the PM has been highly studied for the past decades and an overwhelming amount of data is available in the literature (6-8, 20, 42, 43). Several experimental systems have been used for this goal, both *in vivo* and *in vitro* (14, 16, 19, 27, 42, 43). It is well known that the MA domain of GAG interacts with negatively charged membranes via its N-terminal region, which is myristoylated and contains a cluster of positive charged residues (12, 42). Additionally, MA is capable to interact with oligonucleotides, which may regulate the protein specificity for membranes containing the lipid PIP_2_ (at the inner leaflet of the plasma membrane) (17, 28-30). In this work, we aim to study the interaction of recombinant MA protein with several model membranes (monolayers and bilayers) containing phosphatidylserine (negatively charged lipid) with or without a small amount of PIP_2_ in its composition.

Monolayer techniques are very useful for analyzing the ability of proteins to adsorb/penetrate to different interfaces (37, 44). To do this, molecules are added directly into the bulk of the subphase (36) and changes of the surface tension, which provides information on the interaction process of the molecules with the interface, are registered from the time of injection (36, 41, 44). Based on this, we first studied the ability of MA to spontaneously partition into a clean air-buffer surface allowing us to characterize its tendency to adsorb to interfaces. Figure S2 in the Supporting Information shows an example of these results and table 1 summarizes both the thermodynamic and kinetic parameters obtained from these measures.

**Table 1.** Kinetic and thermodynamic parameters obtained for the MA adsorption to a clean air-buffer interface.

| | $\kappa$ (mN·m<sup>-1</sup>·min<sup>-1</sup>) | $\Pi_{eq}$ (mN·m<sup>-1</sup>) | $\Delta G^{ADS}$ (kcal·mol<sup>-1</sup>) |
| --- | --- | --- | --- |
| **MA\*** | 0.25 ± 0.03 | 21 ± 1 | -8.72 ± 0.09 |
| **myrMA\*** | 0.24 ± 0.03 | 26.4 ± 0.9 | -8.61 ± 0.09 |

\*1,5 µg/mL bulk concentration

From these results it is possible to note that MA has a high propensity to adsorb into the air-water interface, denoted by the high values of ΔG^ADS^, which is in agreement with previous studies on the surface behavior of these proteins (32). On the other hand, the presence of the myristic acid in MA allows the protein to develop slightly higher Π_eq_ values, which corresponds to the maximum pressure difference that can be reached independently of the increase in protein concentration in the subphase (37). Interestingly, this is the only difference observed for the myristoylated protein meaning that, even from a kinetic point of view, this modification does not seem to be influencing the adsorption process at the air-water interface. This result agrees with the hypothesis that the myristic group is probably sequestered in the protein’s core (43) and its presence would not significantly affect its the interfacial adsorption tendency.

For several decades, the monolayer technique has been widely used to study protein interaction with biological membranes (44, 45); however, and surprisingly, this has not been sufficiently exploited for the case of HIV-1 GAG. This system, despite being a simplified version of currently accepted cell membrane models (i.e., bilayers), allows controlling the properties of the interface in a unique way compared to other models. The fact of being able to intervene on the number and type of lipid molecules present in a defined area, offers the possibility of having a total management in the initial conditions of the experiments. The lipid compaction, its phase state, charge density, among several other properties, are not easily controllable variables in other artificial membrane systems and, much less, in cellular models (39, 40, 45-47). Therefore, here we decided to study the association of MA to lipid monolayers by following the lateral pressure variations upon the inclusion of the protein in the subphase. Because of MA is known to bind negatively charged membranes, particularly those containing PIP_2_ (15, 19), three lipid interfaces of interest were used in this work: a neutral one, composed only of POPC; an anionic without PIP_2_ (POPC:DOPS 60:40); and an anionic that includes a small amount of PIP_2_, (POPC:DOPS:PIP_2_ 62:36:2).

Figure 1 shows the surface pressure change due to the protein penetration into the lipid monolayer at different initial lateral density. From these data is possible to obtain a cut-off pressure (Π_C_) that represents the maximal lateral lipid packing from which there is no longer any lateral pressure variations (i.e., protein penetration) after the addition of molecules into the bulk of the subphase (ΔΠ = 0). This parameter is related to the protein’s affinity or penetration capacity into the interface and values higher than the equilibrium pressure (the above-described Π_eq_ values) indicate that the lipids facilitate the adsorption with respect to the air-water surface (36, 41). Table 2 summarizes these parameters for the studied conditions, including the kinetics results corresponding for the adsorption of the protein into lipid monolayers at low lateral packing (low lateral pressure values).

**Table 2.** Kinetic and thermodynamic parameters of the MA partition into lipid monolayer interfaces.

| interface | $\kappa$ (mN·m<sup>-1</sup>·min<sup>-1</sup>) * | | | | |
| --- | --- | --- | --- | --- | --- |
|  | w/o Sel25 |  |  | + Sel25 |  |
|  | PC | PS | PIP<sub>2</sub> | PS | PIP<sub>2</sub> |
| MA** | 0.5 ± 0.1 | 2.0 ± 0.2 | 3.9 ± 0.9 | 1.1 ± 0.2 | 1.2 ± 0.5 |
| myrMA** | 2.3 ± 0.7 | 2.2 ± 0.1 | 3.6 ± 0.4 | 1.6 ± 0.1 | 1.8 ± 0.5 |
| interface | $\Pi_c$ (mN·m<sup>-1</sup>) | | | | |
|  | w/o Sel25 |  |  | + Sel25 |  |
|  | PC | PS | PIP<sub>2</sub> | PS | PIP<sub>2</sub> |
| MA** | 22 ± 2 | 30 ± 2 | 29 ± 3 | 27 ± 1 | 28 ± 2 |
| myrMA** | 26.8 ± 0.5 | 30 ± 1 | 29.9 ± 0.9 | 28 ± 1 | 31 ± 2 |

\*Obtained from adsorption curves at ~10 mN·m<sup>-1</sup> of initial pressure
\*\*1,5 µg/mL of subphase concentration

**Figure 1.**
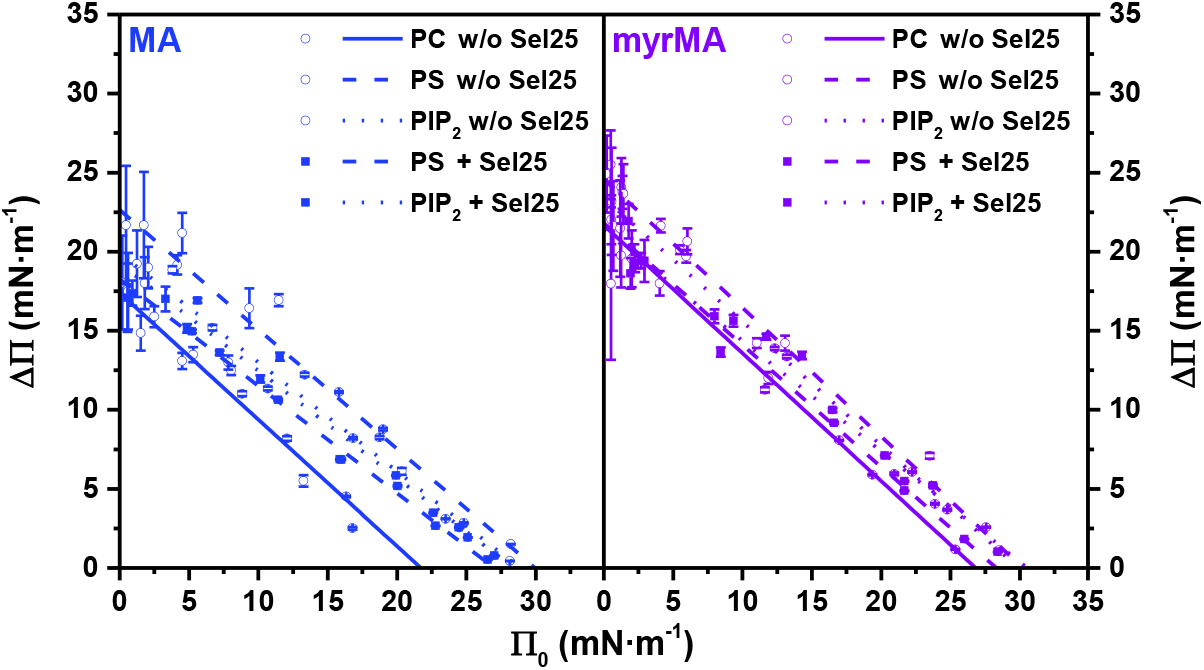
Penetration of MA into different lipid interfaces and the influence of the Sel25 oligonucleotide. The graph shows the change in surface lateral pressure (ΔΠ) upon the adsorption of MA protein into the lipid interfaces at different initial surface pressures (Π_0_). The pre-formed lipid monolayers were composed of POPC (PC, solid line); POPC:DOPS 60:40 (PS, dashed line) and POPC:DOPS:PIP_2_ 62:36:2 (PIP_2_, dotted line). The non-myristoylated (left panel) and the myristoylated (right panel) MA protein were directly injected into the subphase after the monolayer formation and in presence (filled squares) or absence (open circles) of 5-fold amount of Sel25.

From these data it is possible to note that the presence of anionic membranes favors protein adsorption compared to the clean air-water interface, both from a kinetic and thermodynamic point of view. This is deduced from the fact that for the studied conditions κ is always greater with respect to the air-water interface (see table 1) and Π_C_ are greater than Π_eq_. However, given the range of Π_C_ values obtained (around 30 mN/m), it is possible to conclude that MA has a low monolayer penetration capacity (37, 44, 48). This result is expected because this protein is considered to interact superficially with the plasma membrane and might not be significantly inserted into the bilayer (14, 20, 42, 43). In addition, no difference was found due to the presence of the myristic group in the interaction with negatively charged monolayers, indicating the electrostatic nature of the process. However, for neutral lipid films, an increase in both Π_C_ and κ, respect to the clean surface, suggests that hydrophobic interactions might have a role in the absence of charge attraction. This can be in line with previous suggestions that a membrane environment could provoke the exposure of the myristic acid, sequestered when MA is in solution (21, 43, 49), making this modification more important for the membrane anchoring than for the attraction itself.

Interestingly, neither the presence of PIP_2_ nor the preincubation of the protein with the Sel25 RNA appears to have any major influence on the adsorption/penetration process in terms of the achieved Π_C_. This suggests that the presence of this lipid and/or the oligonucleotide, does not seem to have much influence on the final thermodynamic state reached once the protein is at the interface. However, both molecules have a noticeable effect on the kinetics of the process. PIP_2_ by itself increases the protein adsorption velocity (κ values in Table 2), possibly due to a higher charge density. Conversely, Sel25 has a marked effect in reducing this parameter, which is consistent with an effect of electrostatic competition for the basic region of the protein (17, 27, 28). Curiously, the presence of PIP_2_, under these experimental conditions, does not denote a clear ability to reverse this effect of Sel25 on the interaction of MA with membranes. This was not be expected by considering the proposed mechanism of RNA regulation for the interaction of MA with membranes (17). However, given the fact that our experimental system is far from those where this regulation effect is observed, our data suggest that the sole presence of PIP_2_ would not be the only determining factor for MA-membrane specificity and other membranes properties (phase state, electrostatic characteristics) might also be important.

In order to achieve a deeper understanding on the interaction of MA with lipid interfaces, we carried out lateral miscibility studies using Langmuir monolayers composed of defined mixtures of protein and lipids. For these experiments, instead of adding the protein in the subphase, MA is spread directly onto a pre-formed Langmuir lipid monolayer at 0 mN·m^-1^. Once these protein-lipid monolayers are formed, the surface pressure is measured upon variations of the mean molecular area, by mechanical lateral compression, and the lateral pressure-area isotherms are registered (figure 2 A). With this approach, information about the lateral interactions between the monolayer components (i.e., protein and lipids) can be quantitatively obtained (37, 38). Figure 2A shows the compression isotherms of protein-lipid films with different molar fractions of the protein (X_P_) and for various lipid mixtures. From these isotherms it is possible to note, at high proportions of protein, the existence of a first break point (or first discontinuity) that corresponds to the values obtained for monolayers composed only by the protein (arrows and insets in Figure 2A). The fact that these values remain similar for all the mixtures suggests that miscibility is not total under these conditions (37). This indicates that protein-protein interactions could be favored, and a fraction of the molecules is segregated in a different lateral phase with respect to the lipids, maybe due to multimerization.

**Figure 2.**
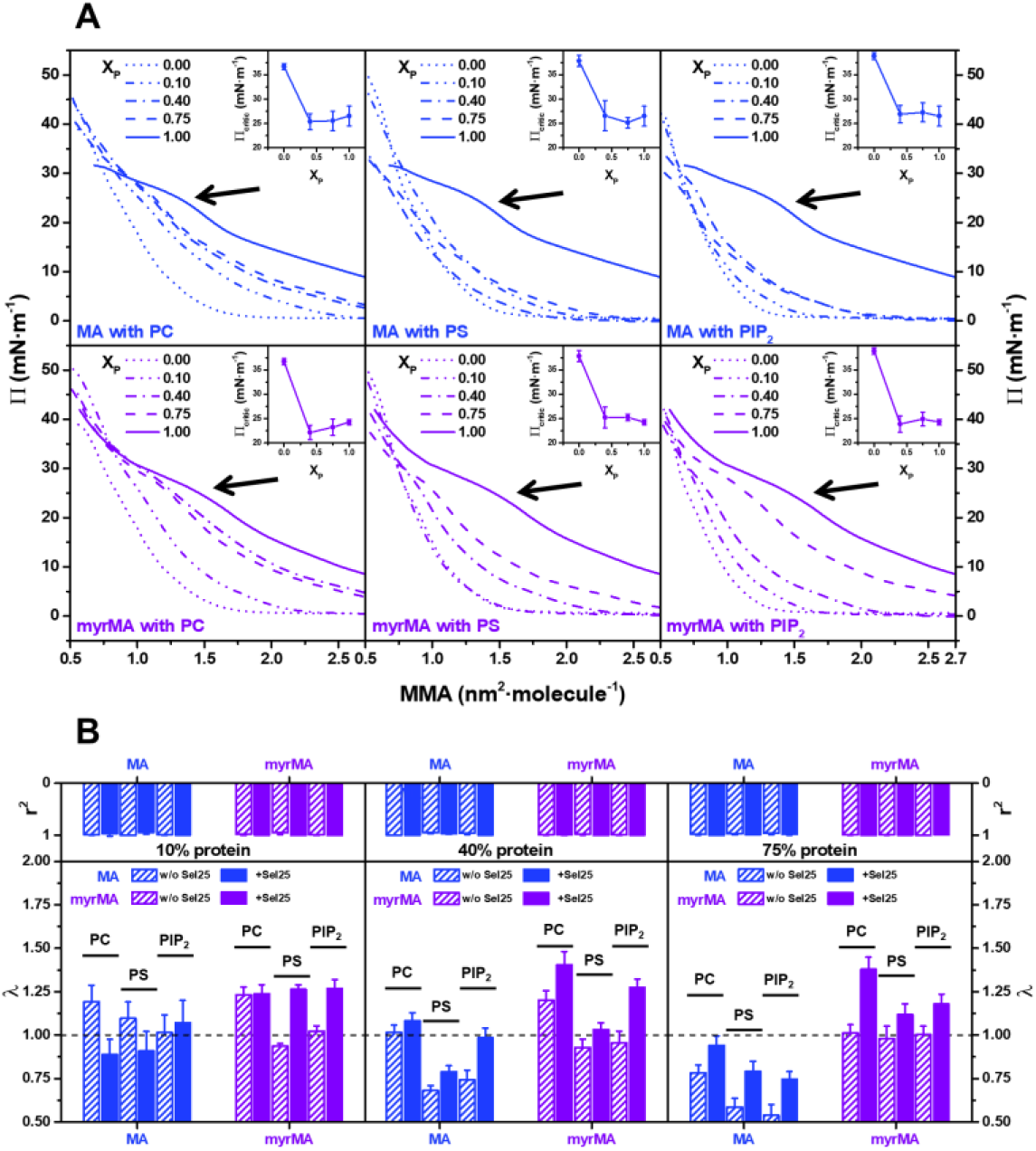
MA lateral miscibility with lipid interfaces. (A) Isothermal variation of the surface pressure (Π) as a function of the mean molecular area (MMA) for monolayers composed of various mixtures of MA (upper plots) or myrMA (lower plots) with lipid interfaces of POPC (PC, left panel), POPC:DOPS 60:40 (PS, center panel) or POPC:DOPS:PIP_2_ 62:36:2 (PIP_2_, right panel). Inserted legend on the top left of each plot denotes the molar fraction of protein in each mixture isotherm and the inset graphs (top right) show the values of the first discontinuity, or Π_critic_, (black arrow) as a function of the molar fraction of protein. (B) Analysis of the Λ-factor for lateral miscibility studies of MA in lipid interfaces. The values of λ (lower panels) and r^2^ (upper panels) are shown for the comparison between the real and ideal compression isotherms of lipid interfaces containing 10% (left), 40% (center) or 75% (right) of proteins. The solid bars indicate the presence of Sel25, and the dotted line indicates the value of λ = 1 as reference of the ideal case.

On the other hand, figure 2B illustrates the results of the obtained Λ-factor for mixtures of MA with lipid interfaces and based on this parameter we carried out the protein-lipid lateral miscibility analysis. The first interesting point of these results consists in the value of r^2^ (top panel on figure 2B), which was always very close to 1 (greater than 0.98 in most cases). This indicates that the differences between the experimental and the ideal isotherms are only given by the value of λ and are independent of Π. On the other hand, the non-myristoylated protein has a clear attraction effect with the lipids in negatively charged interfaces at X_p_ higher than 0.1, since the values of λ are less than 1 and meaning that the experimental molecular areas obtained for the mixed films are lower than those expected for the ideal condition. Contrary, for the case of the myristoylated protein, the values of λ are higher than 1 when mixed with neutral lipids (PC) for X_p_ lesser than 0.4. These data indicate the possibility of repulsive interactions are observed at interfaces composed of neutral lipids; however, in the presence of negative charges, part of this effect could be compensated by attractive forces with anionic lipids, obtaining behaviors closer to the ideal. Additionally, the inclusion of Sel25 shows, in both cases, an increase in the values of λ with respect to the condition without oligonucleotide, particularly for X_p_ higher than 0.1, indicating that the RNA could have a neutralizing effect on some charges of the protein, increasing repulsions at the interface.

### 3.2. MA specificity for PIP_2_ regulated by RNA

In order to study the interaction of the purified proteins with liposomes, we used the approach proposed by Alfadhli et al. of using immobilized protein (27). To achieve this, we took advantage of the fact that the recombinant MA protein contains a hexa-histidine tag located at the carboxyl end, which gives affinity for metal chelates. In this way, it is possible to immobilize the protein on Ni-NTA modified agarose beads so that the amino terminal elements are exposed. These MA-coated beads, visible under the microscope, are incubated with fluorescently labeled liposomes or nucleic acids, allowing detection of their binding (27, 28). In this sense, the quantification of the fluorescent brightness in the region of the beads would be related to the number of liposomes/oligonucleotides that are interacting with the immobilized protein, which allows estimating, in a relative way, the binding efficiency for different experimental conditions. Figure 3 shows a representative example for the binding of DilC_18_-labelled liposomes composed of POPC, POPC:DOPS 60:40 and POPC:DOPS:PIP_2_ 62:36:2 to MA and myrMA coated beads.

**Figure 3.**
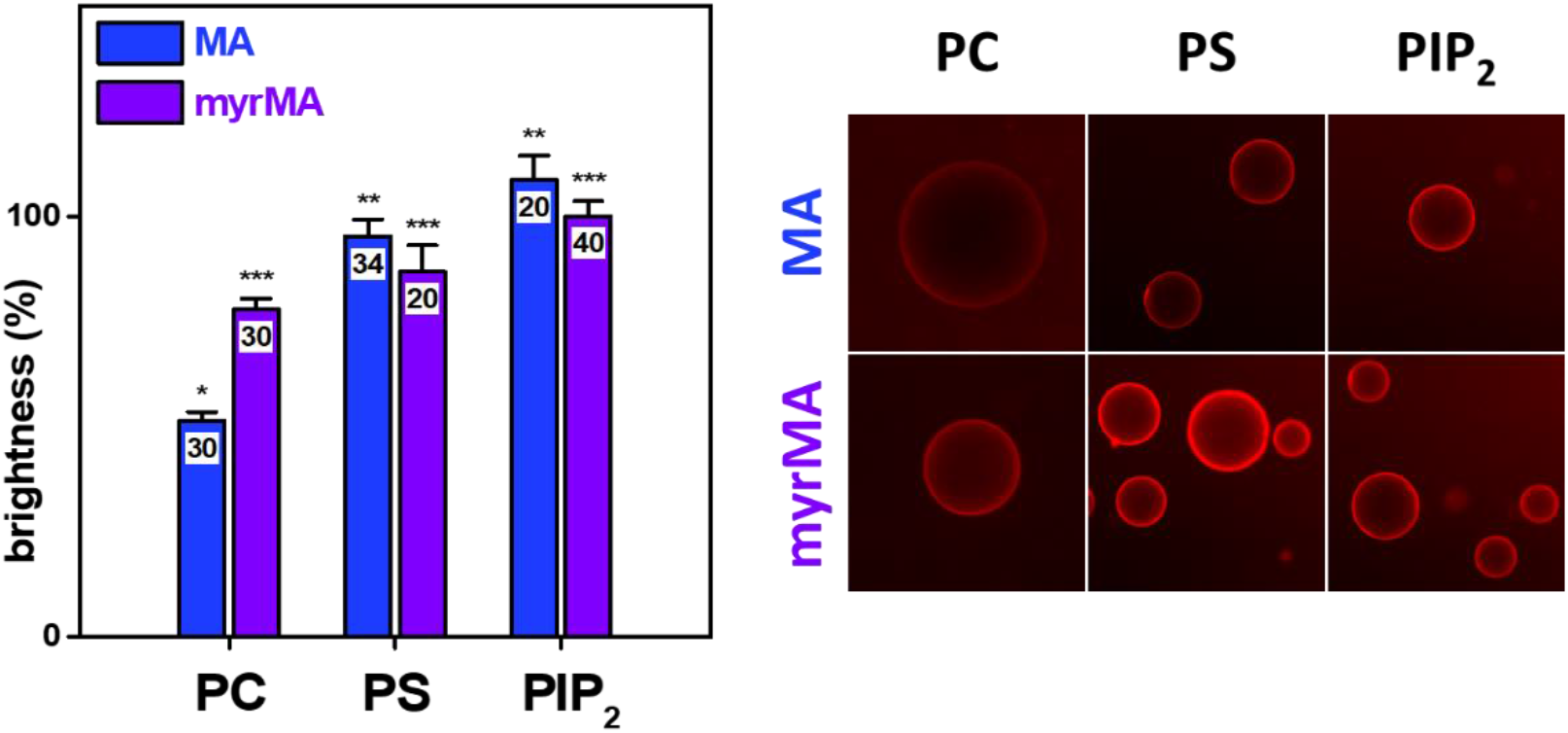
Liposomes binding to MA-coated beads. Quantification of relative brightness in MA (blue) or myrMA (purple) coated beads incubated with POPC (PC), POPC:DOPS 60:40 (PS), and POPC:DOPS:PIP_2_ 62:36:2 (PIP_2_). For the determination of the relative brightness, the case of myrMA incubated with liposomes containing PIP_2_ was used as a reference. The average and its standard error are shown, and the values included inside the bars indicate the total number of beads used for each quantification. The right panel displays some representative images of protein-coated Ni-NTA beads in the presence of red-labeled PC, PS, or PIP_2_ liposomes with 0.5% DilC_18_. The brightness and contrast of the images have been modified to achieve a better visualization; however, the quantification is performed using the original images. Statistical differences represented are obtained by a t-test for p < 0.05.

From figure 3 it is possible to observe that, as expected, the binding of liposomes to the MA-loaded beads is favored when there are negatively charged lipids, while the presence of myristic acid increases the interaction with neutral liposomes, denoting the existence of some hydrophobic contribution, and in line with other results reported previously (15, 20, 27, 42, 43). On the other hand, it is also possible to notice that the inclusion of 2% PIP_2_ in these conditions does not significatively increase the liposomes binding. In previous studies, where an experimental system similar to the one shown here was used, an increase in binding was observed in the presence of PIP_2_ for some conditions (27). However, it is important to clarify that in those studies a greater amount of PIP_2_ (10%) was used and the comparison was made with liposomes of the same percentage of PS, which will have a lower net charge (27). Considering that electrostatic attraction is the main driving force in the binding process, it can be expected that, under conditions of high charge density, the inclusion of a small amount of PIP_2_ will not dramatically increase binding. In this sense, it has already been proposed that this greater sensitivity in the binding of MA to membranes containing PIP_2_ is reduced for liposomes with a higher total surface charge, such as those used here (50). Then, our situation, where PS and PIP_2_ liposome binding is not significatively different, is even more interesting to study the regulation mediated by the interaction with nucleic acids and the possible competition effect of myrMA_21_ peptide.

Several previous studies propose that the MA specificity for PIP_2_ could be regulated via its interaction with nucleic acids (17, 27, 30). Therefore, we assessed the effects of the presence of Sel25 on the binding of liposomes to the protein-coated beads, which is summarized in figure 4. Here it is possible to observe that the liposome association is lower in the presence of Sel25, however, there is a remarkable recovery for liposomes containing PIP_2_, which is higher for the myristoylated case, in accordance with previous results (27, 30). These data indicate that the effect of Sel25 of reducing the binding of PS liposomes to a greater extent than those containing PIP_2_ is still maintained under experimental conditions where the affinity of MA for one or another composition is not different. This suggests that this “molecular competition” between the oligonucleotide and the liposomes cannot be explained solely based on an affinity relationship. In fact, it is possible to appreciate that this reduction of the binding in the presence of Sel25 also occurs -to a lesser extent-for neutral membranes, where a possible electrostatic competition would play a minor role.

**Figure 4.**
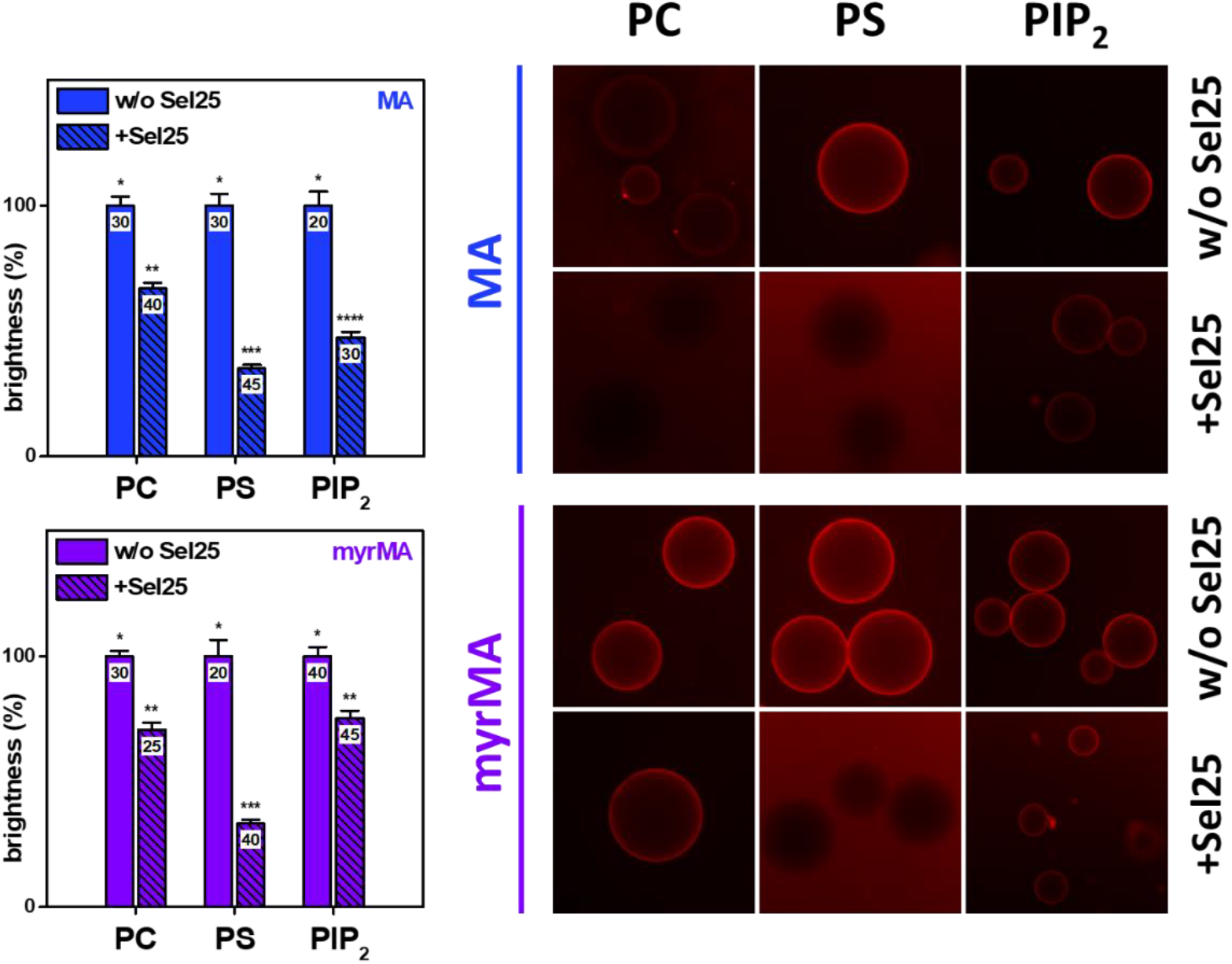
Effect of Sel25 on the liposomes binding to MA-coated beads. Quantification of relative brightness in beads coated with MA (top) or myrMA (bottom) incubated with POPC (PC), POPC:DOPS 60:40 (PS), and POPC:DOPS:PIP_2_ 62:36:2 (PIP_2_) liposomes in the absence or presence of Sel25. For the determination of the relative brightness, the condition without Sel25 was used as a reference in each case. The average and its standard error are shown, and the values included inside the bars indicate the total number of beads used for each quantification. The right panel shows some representative images of protein-coated Ni-NTA beads incubated with red-labeled PC, PS, or PIP_2_ liposomes with 0.5% DilC_18_, in the presence or absence of Sel25. The brightness and contrast of the images have been modified to achieve a better visualization; however, quantification is performed using the original images. Statistical differences represented are obtained by a t-test for p < 0.05.

### 3.3. Competition of myrMA_21_ peptide for the binding of MA to RNA and liposomes

One of the advantages of the above-described experimental system consists in its use to study the effect that other molecules could present on the union of MA with liposomes and/or nucleic acids. Similar systems have already been used for this purpose, including studies of inhibition of MA-RNA interaction mediated by various compounds (27, 28). Thus, we decided to investigate whether the MA interactions with oligonucleotides and/or lipids can be affected by the presence of the N-terminal peptide, whose binding to nucleic acids and membranes has been the subject of study in our previous works (32, 33). In this sense, figure 5 shows the effects of myrMA_21_ on FITC-Sel25 binding to MA protein. As can be clearly seen in the upper images, in the absence of peptide there is an evident binding of Sel25 to the protein-coated beads, a condition that is used as a reference for quantification.

**Figure 5.**
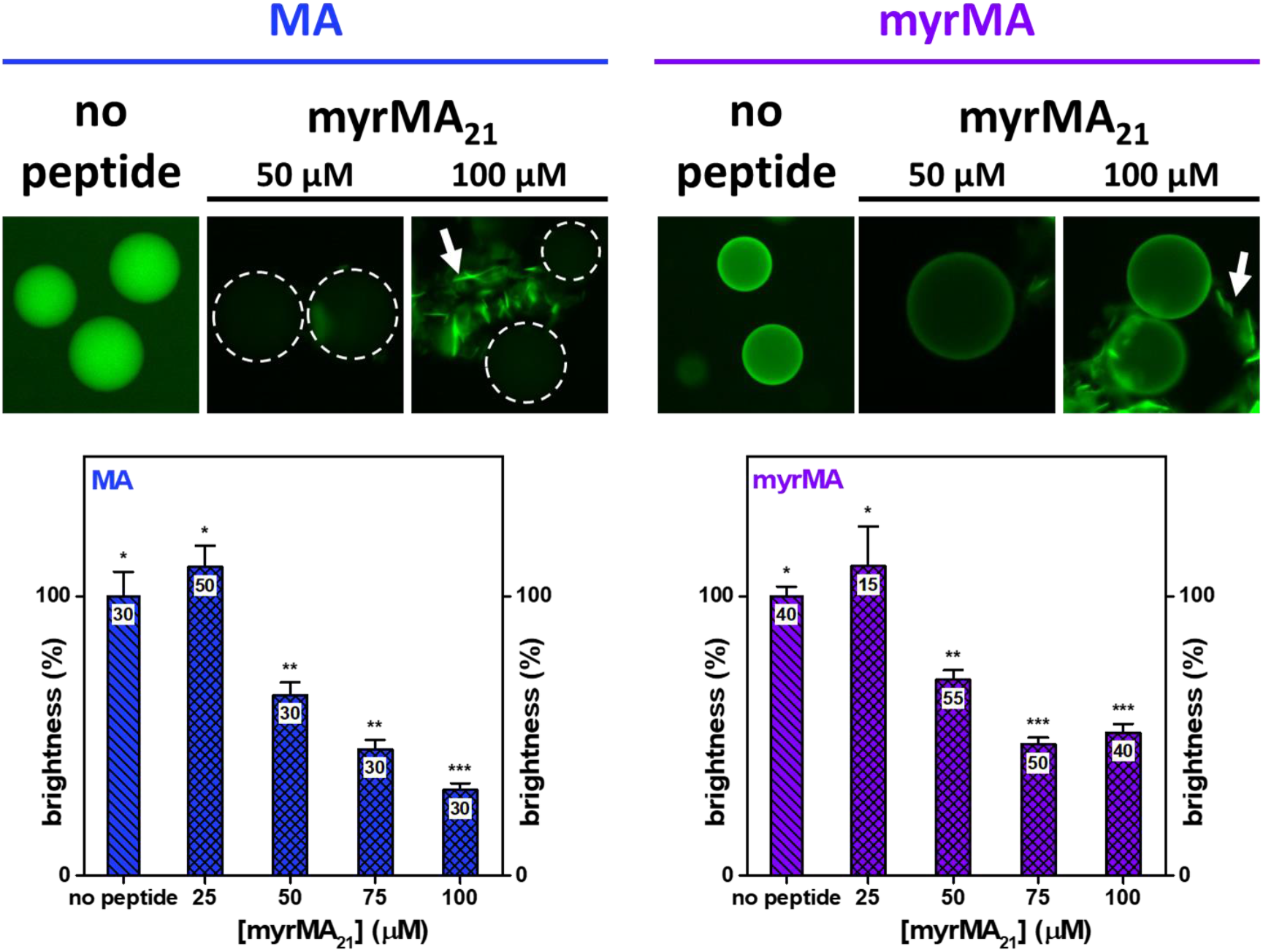
Effect of myrMA_21_ peptide on Sel25 oligonucleotide binding to MA-coated beads. Competition of myrMA_21_ for Sel25 binding to beads coated with MA (blue) or myrMA (purple). Representative images of protein-coated Ni-NTA beads incubated with a FITC-labeled Sel25 variant, in the absence or presence of myrMA_21_ peptide, are shown at the top of the figure. The white arrow indicates the appearance of supramolecular aggregates in the presence of myrMA_21_. The dotted circles roughly distinguish the border of the beads on cases showing low RNA binding. The variation in relative brightness for increasing amounts of myrMA_21_ is shown at the bottom. For the determination of the relative brightness, the condition without peptide was used as reference in each case. The average and its standard error are shown, and the values included inside the bars indicate the total number of beads used for each quantification. The brightness and contrast of the images have been modified to achieve a better visualization; however, quantification is performed using the original images. Statistical differences represented are obtained by a t-test for p < 0.05.

From these results, it is possible to notice that increasing concentrations of peptide decreases the binding of the oligonucleotide to MA. These data demonstrate that the peptide binds to Sel25 and is being capable to compete with the protein for the union of the oligonucleotide. In a previous work, we proposed that this peptide massively interacts with oligonucleotides, which is favored by the formation of peptide/Sel25 aggregates in solution (32). These microscopy results are new compelling evidence of this possibility, since it is possible to observe the existence of large aggregates at high concentrations of myrMA_21_ that bind FITC-Sel25 (arrows in the images of figure 5). Interestingly, the peptide competition for the oligonucleotide is lower for the myristoylated variant of the protein. This indicates that the presence of the myristic acid in the amino terminal region of MA has some effect on the RNA interaction, promoting a better retention of the oligonucleotide. In the context of high protein concentration provided by the immobilization conditions, the conformational state of the myristic group could contribute to achieve a better exposure of the residues and a higher effective charge density on the N-terminal region, leading to a tighter interaction with the RNA.

In line with the above-described results, we decided to test whether the peptide has any effect on the binding of liposomes to Ni-NTA beads coated with MA protein, which is illustrated in figure 6. From this, it is possible to see a clear effect of myrMA_21_ peptide in reducing the interaction of MA with PS liposomes. This effect is less pronounced for beads coated with myristoylated protein, in agreement with our previous observations that the myristic group increases protein binding to membranes. Remarkably, in the presence of liposomes containing 2% PIP_2_, this competition effect is lost, since no significant differences are observed in the presence of myrMA_21_. We showed before that the partition of myrMA_21_ to membranes is not influenced by the presence of PIP_2_ (33), meaning that the dissimilarities observed here are not due to less peptide binding to PIP_2_ liposomes. This result is extremely interesting considering that, in this experimental context, the affinity of the MA protein for charged liposomes is not very different in the presence of PIP_2_ (figure 3). In this sense, a possible explanation could be that the permanence of some peptide fractions bound to the membrane (regardless of PS or PIP_2_) causes changes in the electrostatic properties of the liposomes. This could lead to a reduction in surface charge density, resulting in an accentuation of the MA sensitivity by membranes containing a small amount of PIP_2_, as has been proposed before (50).

**Figure 6.**
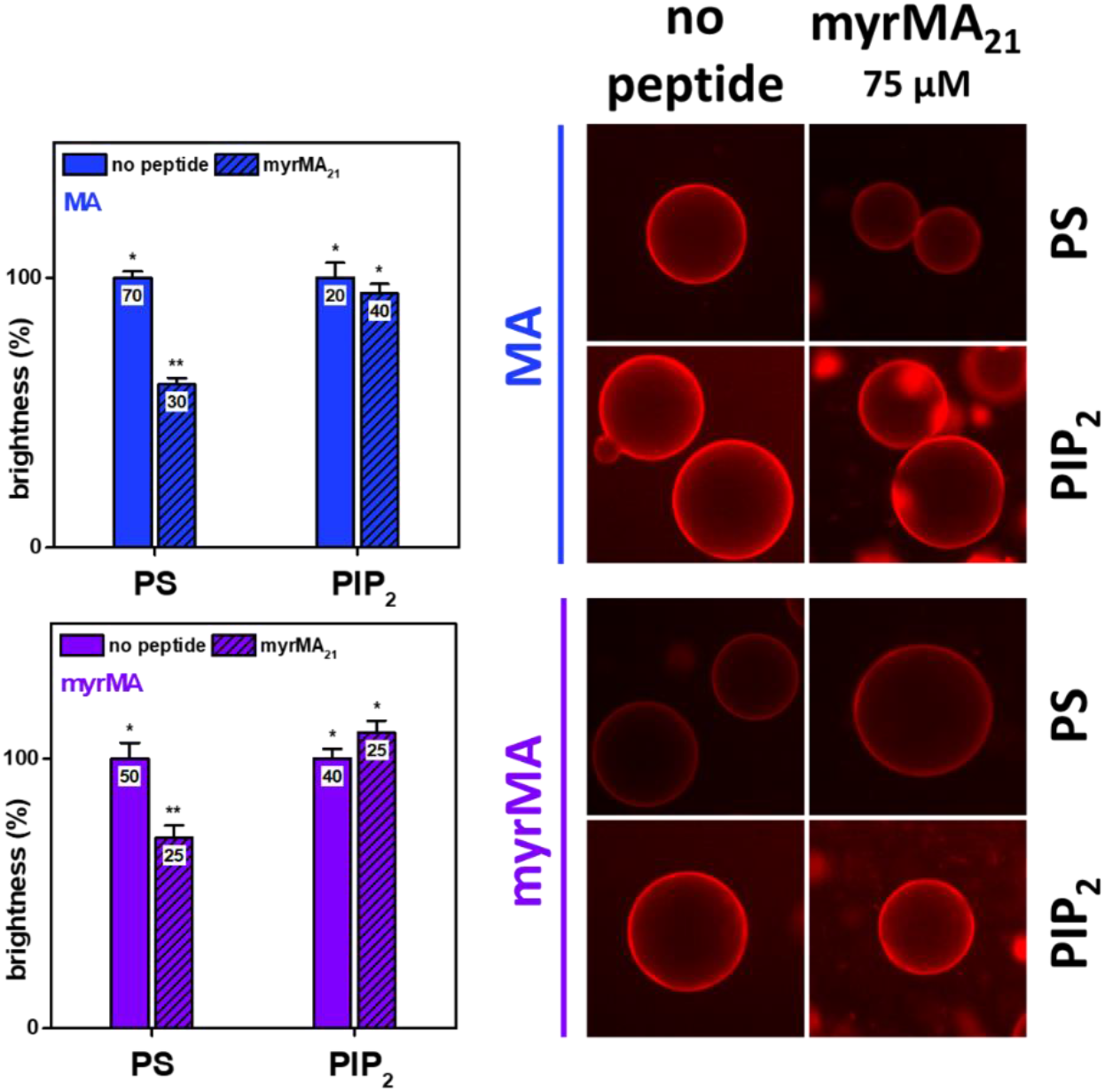
Effect of myrMA_21_ peptide on liposome binding to MA-coated beads. Quantification of relative brightness in beads coated with MA (top) or myrMA (bottom) incubated with POPC:DOPS 60:40 (PS) or POPC:DOPS:PIP_2_ 62:36:2 (PIP_2_) liposomes in absence or presence of myrMA_21_. For the determination of the relative brightness, the condition without peptide was used as reference in each case. The average and its standard error are shown, and the values included inside the bars indicate the total number of beads used for each quantification. The right panel shows some representative images of protein-coated Ni-NTA beads in the presence of red-labeled PS or PIP_2_ liposomes with 0.5% DilC_18_, in the presence or absence of 75 μM of myrMA_21_. The brightness and contrast of the images have been modified to achieve a better visualization; however, quantification is performed using the original images. Statistical differences represented are obtained by a t-test for p < 0.05.

### 3.4. RNA regulation for the MA specificity to PIP_2_ cannot be overcome by myrMA_21_

On the other hand, knowing that myrMA_21_ can strongly compete for the interaction of the protein with Sel25, we were interested in investigating if the presence of this peptide can overcome the effect of the oligonucleotide in preventing binding with non-PIP_2_ membranes. For this, we seize the opportunity that this experimental system allows us to observe more than one component simultaneously. In this sense, experiments were carried out in the presence of FITC labelled Sel25 (green) and negatively charged liposomes (with or without PIP_2_) labeled with DilC_18_ (red). Figure 7 summarizes the results of these tests.

**Figure 7.**
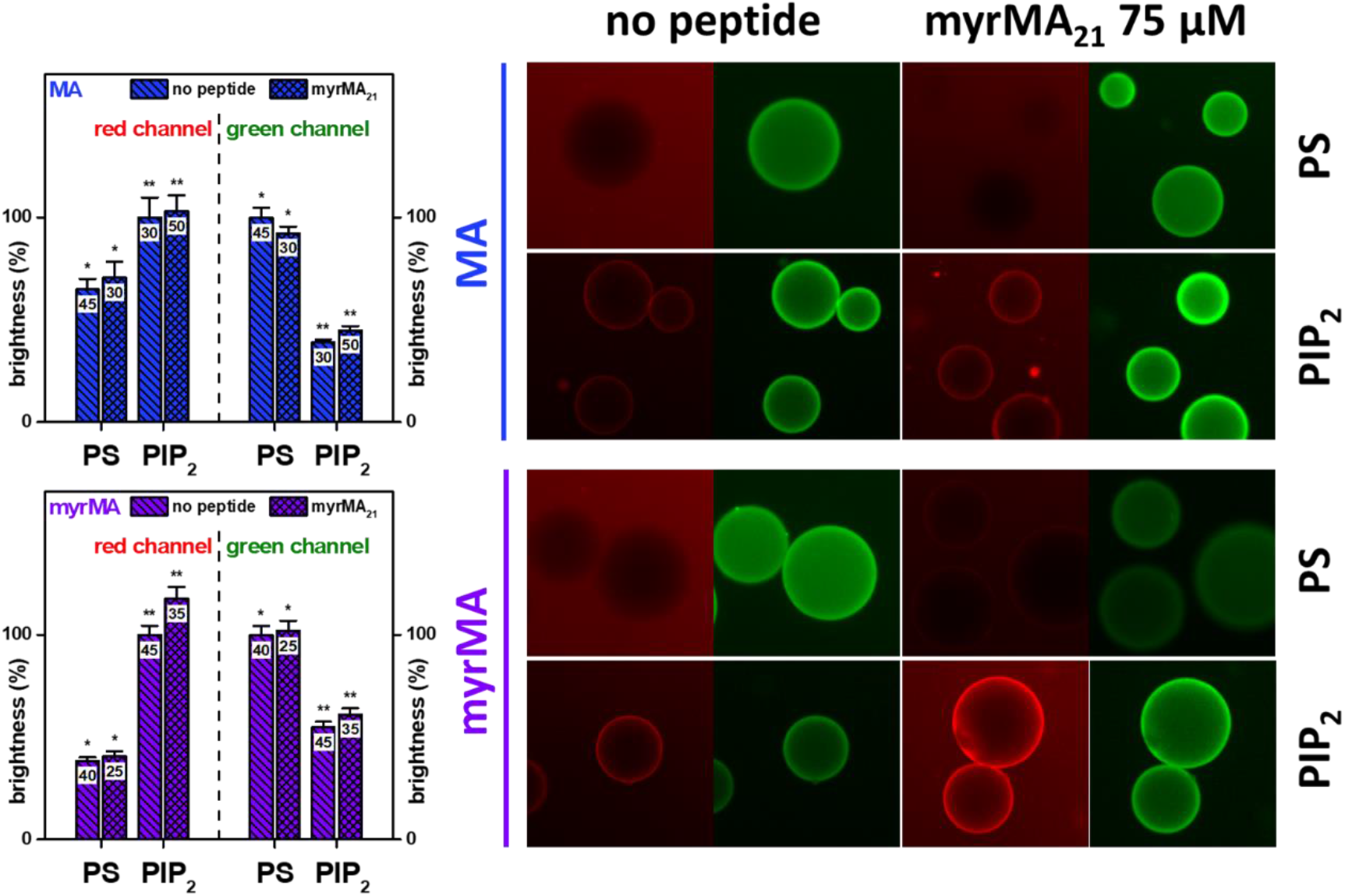
Effect of myrMA_21_ peptide on the Sel25 and liposomes binding to MA-coated beads. Quantification of relative brightness in beads coated with MA (top) or myrMA (bottom) incubated with FITC-Sel25 and POPC:DOPS 60:40 (PS) or POPC:DOPS:PIP_2_ 62:36:2 (PIP_2_), in absence or presence of myrMA_21_. The bars on the left denote the quantification in the red channel (liposomes) while the bars on the right denote the quantification in the green channel (oligonucleotides). For the determination of the relative brightness, the brightest condition without peptide was used as a reference in each case. The average and its standard error are shown, and the values included inside the bars indicate the total number of beads used for each quantification. The right panel shows some representative images of protein-coated Ni-NTA beads incubated with FITC-Sel25 and red-labeled PS or PIP_2_ liposomes with 0.5% DilC_18_, in the presence or absence of 75 μM of myrMA_21_. The red channel is shown on the left and the green channel on the right for the same image. The brightness and contrast of the images have been modified to achieve a better visualization; however, quantification is performed using the original images. Statistical differences represented are obtained by a t-test for p < 0.05.

From these results it is possible to highlight that, in the absence of the myrMA_21_ peptide, there is a greater binding to liposomes containing PIP_2_ compared to those containing only PS (red channel). This is in line with what was discussed above (figure 6), where it was already observed that the difference in binding to PS with respect to PIP_2_ is more notable in the presence of Sel25. Furthermore, it is possible to see that this effect is accompanied by a reduction in the binding of Sel25 to the beads (green channel). Basically, under the conditions where greater liposome binding is observed, there is less oligonucleotide binding and vice versa. This can be seen in the images and quantification in figure 7, where a loss of membrane binding for PS liposomes is observed, which is recovered for PIP_2_ together with a decrease in Sel25 binding. Unexpectedly, in the presence of myrMA_21_ no significant differences were observed compared to the case without peptide. This suggests that, although this peptide is capable of strongly reduce the binding of MA to Sel25, in the presence of liposomes this competition effect is lost. This result could indicate that the peptide can be marginally bound to both liposomes and RNA, leading to a possible recovery of the MA-RNA interaction.

Here and in previous works (32, 33) we have shown how remarkable is the interaction of myrMA_21_ with oligonucleotides and its consequences, including the impossibility of forming monolayers at the water-air interface or the total loss of electrophoretic mobility (32). These effects have been attributed to the high propensity of the peptide to form supramolecular aggregates that could be strongly influencing the interaction with oligonucleotides. In this way, it could be expected that this binding between the peptide and RNA reduces the “chaperone effect” of the latter, which would lead to a greater recovery of the MA-membrane interaction even in conditions where there is no PIP_2_ present. However, this does not occur under the experimental conditions used. Considering the physicochemical characteristics of this peptide (a polybasic molecule capable of forming supramolecular particles with a high charge density and that strongly interacts with oligonucleotides), it is remarkable how its presence is not enough to reverse the regulatory capacity of the association of MA to membranes given by its interaction with nucleic acids.

## 4. Discussion

The assembly of new HIV viral particles is a process that occurs at the internal interface of the cell surface, where the MA domain of GAG plays an important role in establishing this first interaction with membranes (13, 42, 43). In this work we study for the first time the interaction of MA with lipid interfaces using the Langmuir monolayer approach. The use of monolayer techniques has its own advantages that are invaluable, including the possibility of studying interactions with interfaces in conditions of high protein density, a context much closer to cell membranes and that is not easily achievable with other systems (37-40). In this sense, figure 8 shows a schematic summary of the interpretations of the results obtained for the interaction of MA with both, monolayer and bilayer membranes systems.

**Figure 8.**
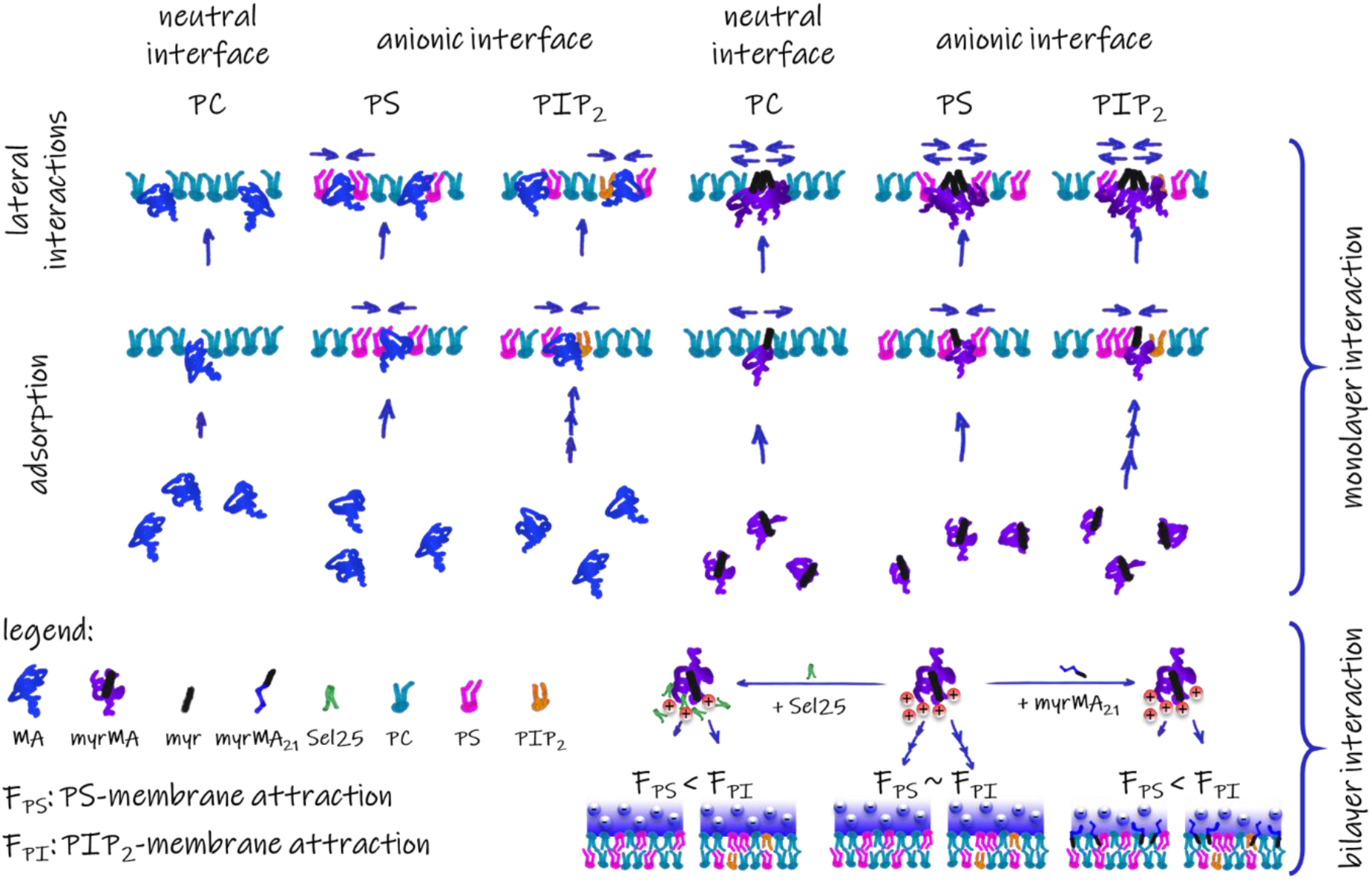
Summary model of the interaction of MA with monolayer and bilayer membranes model. The drawing consists of a schematic representation of the results described in the text. The bottom-right part of the model illustrates the regulation of MA specificity for PIP_2_ driven by both Sel25 and myrMA_21_. This finely tuned modulation of the dielectric properties will affect the global electrostatic attraction, making more prominent the specific contribution of the PIP_2_ lipid. The illustrated molecules are indicated in the legend. myr: myristic acid.

The first important aspect observed here is that anionic interfaces are more favorable for the interaction with MA, both kinetically and thermodynamically, which results in more attractive lateral interactions. This feature agrees with other previous observations (15, 27) and is perfectly explainable based on the physicochemical properties of the protein, whose net charge is positive in all used conditions. Additionally, the myrMA protein shows higher capacity to penetrate neutral membranes. This leads to greater repulsive interactions since more penetration leads to more presence of the polar groups of the amino acid side chains. In this way, in interfaces that also contain anionic lipids, there could be a compensation effect between attractive and repulsive lateral interactions with myrMA. Interestingly, this compensation effect is even greater when there is a greater proportion of protein in the interface. This fact can be explained in a context where lateral protein-protein interactions are favored. These data are in agreement with the case of myrMA, since it is known that membrane environments have the capacity to modulate the exposure of the myristic group of the protein (21, 49), which is related to the propensity to form multimers (34).

On the other hand, it is interesting to note that the presence of PIP_2_ increases the kinetics of adsorption of the protein to the interface, but without having any appreciable influence on other aspects, such as the penetration capacity or lateral protein-protein interactions. In addition, the effects of reducing the interactions of MA with the interface observed in the presence of Sel25 are not reversed by the presence of this inositol. This proposed effect of PIP_2_ to reduce the interaction of MA with nucleic acids has been consistently observed in several studies using other experimental systems (19, 27), including here using vesicles (figure 4). The fact that this phenomenon is not observed in the monolayer system, where PIP_2_ could be found in conditions far from the case of a liposome, suggests that the sole presence of this lipid is not necessarily the main cause for this differential interaction. Therefore, its environment and the physicochemical properties of the membrane could also be important.

As said, we wanted to study possible factors that modulate the specificity of MA for PIP_2_-containing membranes. One, already discussed in several works, is the charge density of the membrane (18, 43, 50). For example, it is known that the inclusion of cholesterol on bilayer systems leads to better union when PIP_2_ is present, which is explained by higher charge density due to a clustering effect (27, 50). Therefore, a regulation of the electrical properties of the protein-membrane association might be crucial. Here we used a model system in where PS and PS/PIP_2_ membranes do not display major differences in binding MA. This lack of specificity can be explained by the fact that these compositions are highly charged, leading to a strong electrostatic attraction that will probably overcome any specific contribution from the PIP_2_ lipid (present at low proportions in our membrane systems). In this sense, we observed that, even in this condition, the presence of RNA increases the protein selectivity for PIP_2_-containing membranes. Consequently, a simple linear relationship of affinities is not enough to explain the results. This have been proposed before by considering that the MA affinity to PIP_2_ is higher than to RNA and the affinity of the protein to RNA is higher than to PS, a situation that will imply that the affinity for PS-PIP_2_ containing membranes is higher than the affinity for those composed of PS, which is not true in our system. Considering this, we propose a more complex mechanism that ensures the MA specificity for the PM, consisting of a finely tuned effect on the electrostatic properties of both membrane and protein’s environment. The bottom-left part of figure 8 attempts to illustrate this proposal and explain our results.

In this sense, the high charge density is a condition where the driving force for the protein interaction is mainly electrostatic and do not discriminate if a small amount of PIP_2_ if present (middle part on the schema). Then, when RNA is included (left part on the schema), the dielectric properties of the protein environment change, resulting in a global reduction of the electrostatic attraction and provoking that the specific interaction for PIP_2_ is now more important. This hypothesis agrees with previous observations that oligonucleotide interaction increases the dielectric constant of the HBR region, which was assessed by fluorescence studies for the N-terminal peptide (33) and corroborated here for the full MA protein (Supporting Information section S3). Additionally, the same argument can explain our results for the myrMA_21_ effect on the liposome binding to MA (figure 6), where, surprisingly, the peptide also increases selectivity for PIP_2_-containing membranes (i.e., it reduces the protein association with PS membranes but not with PIP_2_ membranes). For this case, the peptide might also be modulating electrical properties of the system, this time in the membrane side (figure 8, right part). This also correlates with our previous observations that the peptide can vary certain membranes properties, like its electrophoretic mobility and its ζ-potential (33). In fact, this way for explaining protein-membrane interaction is more accurate if we bear in mind the importance of a correct regulation of the electrostatic properties of a living cell system. Therefore, this new proposed scenario could open the possibility for better understand the regulation of the specificity of MA for PIP_2_-containing membranes by specifically studying macromolecular properties of the system instead of considering only molecular interactions in a simple ligand-receptor view.

## Abbreviations

PM: plasma membrane;
MA: matrix domain;
HBR: highly basic region;
FITC: fluorescein isothiocyanate;
PIP_2_: phosphatidylinositol-(4,5)-bisphosphate;
PC: phosphatidylcholine;
PS: phosphatidylserine.

## Acknowledgments

This work is supported by CONICET and FONCyT (PICT 2018-3204). L.B.P.S. holds a postdoc fellowship from CONICET and E.E.A is a Career Member of CONICET. We are also grateful to all the members of the biophysical area at CIQUIBIC-DQB for helpful discussions.

## Author contributions

The experimental work was performed by L.B.P.S. The project was designed by all authors and directed by E.E.A. The manuscript was written through contributions of all authors. All authors have given approval to the final version of the manuscript.

## SUPPORTING INFORMATION

### S1. Comparing Π–MMA isotherms with the Λ-factor

Performing a comparison between compression isotherms (Π–MMA) of monolayers is an important step to extract information about the characteristics of the molecules forming the film. For this evaluation, we use the previously introduced Λ-factor (1), whose idea is to have a way of quantitatively compare isotherms in an integral way and independent of an arbitrary data point value. In this sense, given two isotherms to compare (one of “interest” and other as “reference”), Λ is defined as the pair of real numbers λ and r^2^:

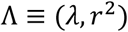

Where λ represents a “distance” factor between the MMA of the two compared isotherms and r^2^ denotes the degree of “similarity” between the shape of the curves throughout the range of pressures common to both isotherms.

Figure S1 illustrates the procedure used to calculate these parameters. To do this, first the range of lateral pressure values that are common to both isotherms (reference and interest) is determined and a nonlinear regression algorithm is used to find the value of λ that minimizes the difference of squares between the values of the MMA for the isotherm of interest and those calculated for each pressure value according to:

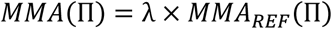

Then, the value of the coefficient of determination (r^2^) is obtained from the comparison between the resulting curve and the curve of interest.

**Figure S1.**
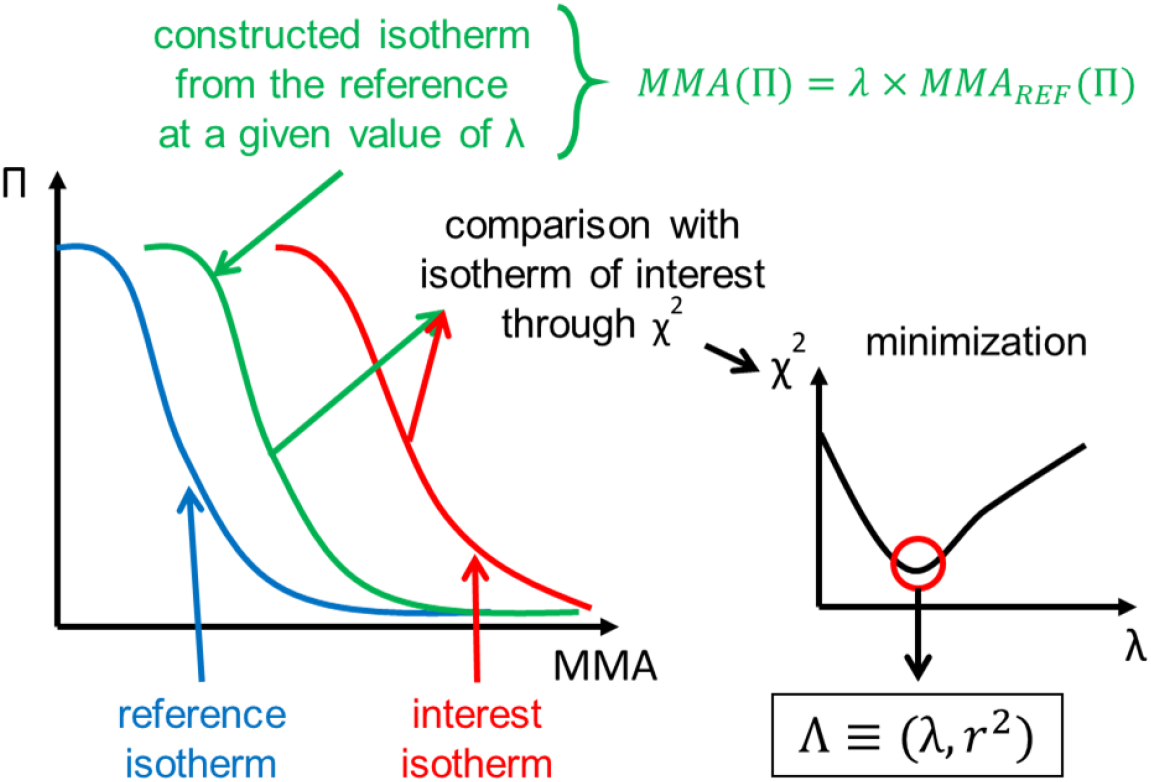
Schema for the algorithm to calculate Λ.

In this sense, when two identical isotherms are compared, both λ and r^2^ are expected to be equal to 1. Now, if the curves are offset from each other in terms of the MMA, but are very similar in shape, then r^2^ would have values close to 1 while the value of λ would quantitatively indicate the degree of displacement, either towards smaller (λ < 1) or larger (λ > 1) molecular areas. For example, one possible reason for such deviations is a change in the number of molecules on the surface, causing an underestimation or overestimation of the molecular area and thus a proportional shift of the curves. On the other hand, if the isotherms are not similar, then r^2^ will have values less than 1 due to differences in the shape of the compared curves, which may indicate structural changes, lateral rearrangements, or molecular transitions.

This Λ-factor is also useful for the analysis of mixtures. For this, the comparison is made between the isotherm obtained experimentally, using as reference those built for the ideal behavior from the pure elements. In this way, the values of λ would indicate the relationship between the real behavior with respect to the ideal, so that its deviations from the unit would correspond to the existence of repulsive (λ > 1) or attractive (λ < 1) interactions. In addition, this analysis allows the inclusion of a parameter (r^2^) that quantifies the similarity between the ideal and real isotherms of the mixtures.

### S2. MA adsorption to the air-water surface

**Figure S2.**
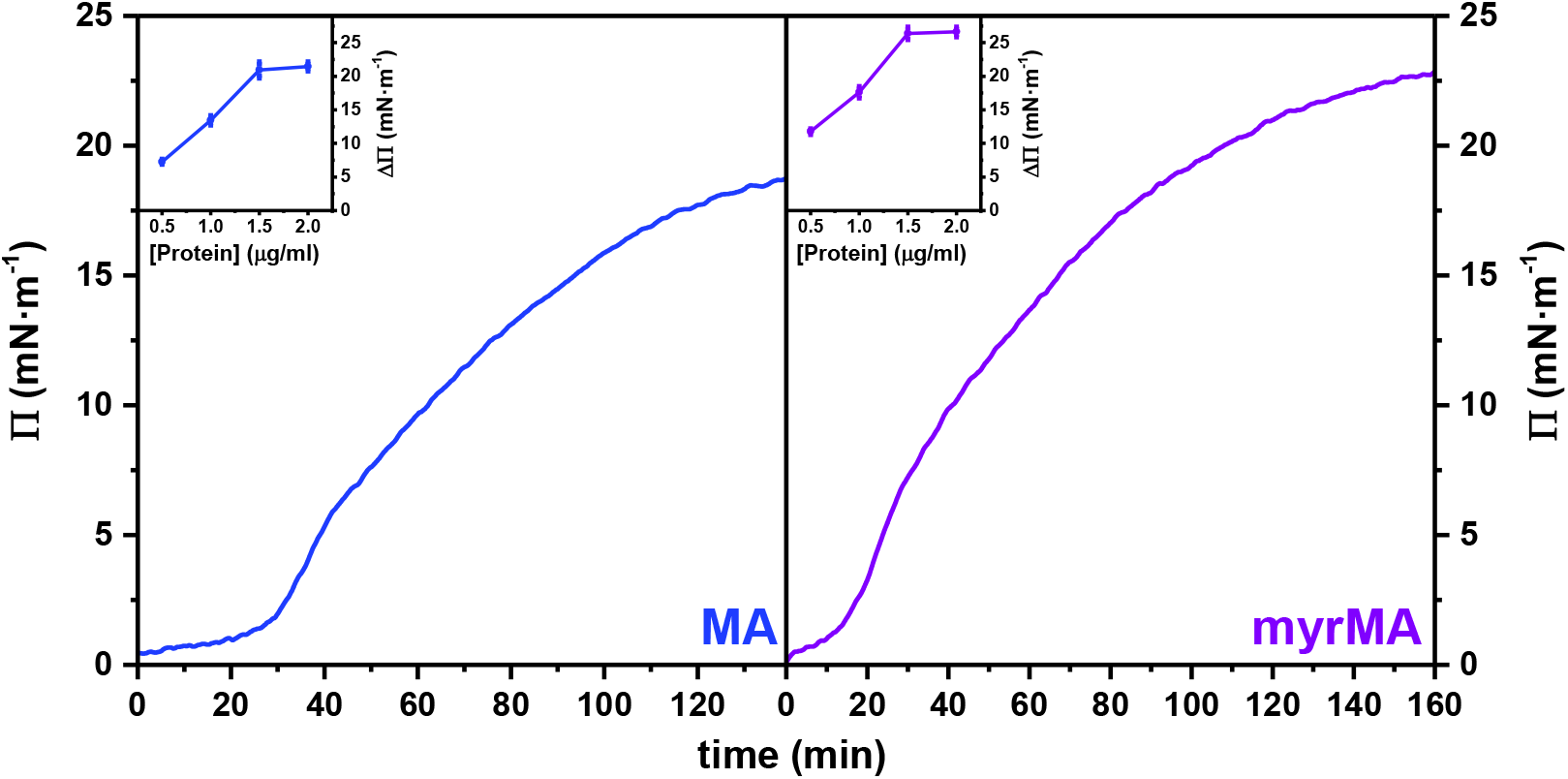
MA adsorption kinetics at the water-air interface. The main plots show the time variation of lateral pressure from the inclusion in the subphase of 1.5 μg/mL of MA (blue) and myrMA (purple). The graphs inserted in each case represent the maximum variation of surface pressure detected as a function of the concentration of protein included in the subphase.

### S3. Oligonucleotide presence affects the polarity of the MA HBR region

To analyze the polarity in the local environment of the proteins, the fluorescent properties of the W residues in the presence or absence of oligonucleotide were determined as previously described (2). For this, we use a Cary Eclipse spectrofluorometer (USA) with a cell of 3 mm optical path length and using slits for excitation and emission of 5 nm each. Detection was performed with a photomultiplier set to 600 V and a 1 nm step was used at a scan speed of 600 nm·min^-1^. All measured samples were prepared in 20 mM Tris-HCl (pH 7.4) with 150 mM NaCl. For the estimation of the dielectric constant from the fluorescence of tryptophan, a 10 μM solution of N-acetyl-L-tryptophan amide, or NATA, prepared in different solvents with known dielectric constant values, was used, as described elsewhere (2). These were: water (80), dioxane 5% (78), dioxane 15% (69), dioxane 25% (57), dioxane 35% (46), dioxane 45% (36), dioxane 55% (27), absolute ethanol (24.5), butanol (17.5). Figure S3 and table S1 summarize the dielectric constant values estimated for MA and myrMA, as well as those corresponding to the peptide myrMA_21_, extracted from Socas & Ambroggio 2020 (2).

**Figure S3.**
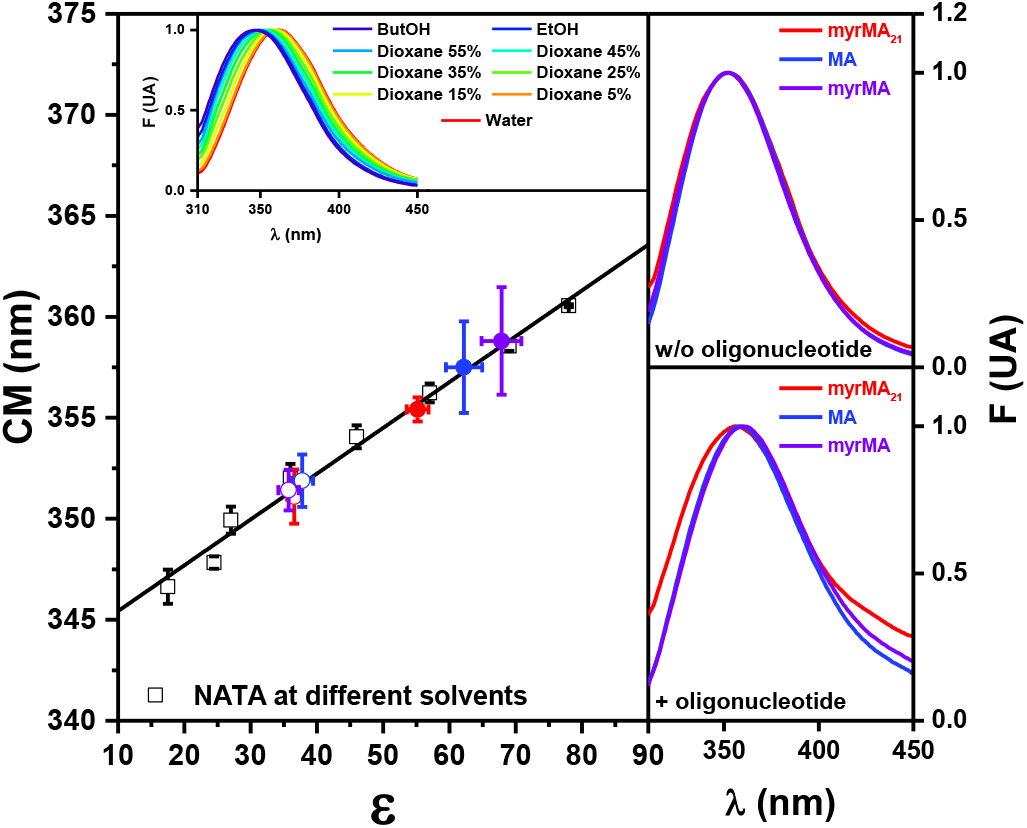
Estimation of the dielectric constant in the local environment of MA protein and myrMA_21_ peptide. (A) Center of mass of the fluorescence emission spectrum of NATA in different solvents with known ε (black squares). A linear model was used to estimate ε. Circles represent the case of MA (blue), myrMA (purple) and myrMA_21_ (red); filled symbols correspond to the presence of 50 μM of oligonucleotide and open symbols correspond to its absence. Top inset in A: emission spectra of NATA in different solvents. (B) and (C) Emission spectra of MA (blue), myrMA (purple) and myrMA21 (red), in the absence (B) or presence (C) of oligonucleotide. Data for the calibration curve and for the peptide myrMA_21_ are extracted from Socas & Ambroggio 2020 (2).

**Table S1.** Spectral center of mass and ε values estimated for the tryptophan environment of the MA protein and the myrMA_21_ peptide.

|  | MA | myrMA | myrMA<sub>21</sub> |
| --- | --- | --- | --- |
| **w/o oligonucleotide** |  |  |  |
| **CM (nm)** | 352 ± 1 | 351 ± 1 | 351 ± 1 |
| **$\epsilon$** | 38 ± 2 | 36 ± 2 | 37 ± 1 |
| **+ oligonucleotide** |  |  |  |
| **CM (nm)** | 358 ± 2 | 359 ± 3 | 355.4 ± 0.6 |
| **$\epsilon$** | 62 ± 3 | 68 ± 3 | 55 ± 2 |

As can be seen from the data, the values obtained for MA and myrMA are very similar to each other, depicting the existence of a more polar environment in the presence of the oligonucleotide. This indicates, on one hand, that the presence of the myristic group does not make any difference in the emission of fluorescence, which is consistent with the fact that this post-translational modification does not entail drastic changes in the structure of the protein that can be reflected in the environment of its fluorescent residues. On the other hand, the variation in fluorescence emission due to the presence of oligonucleotides is indicative that its presence affects the local polarity of the region near the HBR, where one of the tryptophan residues of the protein is located, as was already described for the peptide myrMA_21_ (2).

